# Dose response comparison of Nipah virus strains Malaysia and Bangladesh in hamsters exposed by the intranasal or intraperitoneal route

**DOI:** 10.1101/2025.01.26.634958

**Authors:** Sara C. Johnston, Ju Qiu, Sarah LW Norris, Rekha Panchal, Elizabeth M. Punger, Melissa Teague, Joshua L. Moore, David N. Dyer, Ondraya M. Frick, Stephen C. Stevens, Jimmy Fiallos, Harold L. Mills, Eugene L. Blue, Willie B. Sifford, Trevor McCarson, Amanda Casselman, Jonathan D. Latty, Kathleen E. Huie, Ryan Adolphi, Aysegul Nalca

## Abstract

Nipah virus, a zoonotic pathogen, can cause debilitating disease and death in humans. Currently, countermeasures are limited, with several in various stages of testing but none yet FDA-approved for human use. Evaluation of countermeasure candidates requires safety testing in humans, as well as efficacy testing against lethal challenge in animal models. Herein, we describe the characterization and comparison of the intraperitoneal and intranasal Syrian golden hamster models for Nipah virus strains Malaysia and Bangladesh. Overall, the intraperitoneal route of exposure resulted in a more consistent lethal outcome, regardless of virus strain. Therefore, the IP model was subsequently used to evaluate the use of Favipiravir as a potential positive control for future studies investigating NiV countermeasures. In contrast to prior reported results regarding Favipiravir in Nipah virus-infected hamsters, Favipiravir was only fifty percent effective at preventing death following lethal challenge, regardless of Nipah virus strain. The data suggest that Favipiravir is only partially protective against Nipah virus in hamsters, and, thus, would likely not be an ideal candidate as a positive control in future efficacy studies.

## Introduction

Nipah virus (NiV) is a zoonotic pathogen capable of causing debilitating disease in humans, with a case fatality rate (CFR) between 40-75% according to recent estimates by the World Health Organization (https://www.who.int/news-room/fact-sheets/detail/nipah-virus). There are two distinct stains of NiV, Malaysia and Bangladesh, which differ in their presentation, pathogenicity, and transmissibility. NiV strain Malaysia (NiV_M_) was identified in 1999 following an outbreak in Malaysia and Singapore associated with infected pigs [1]. Since 1999, outbreaks associated with this strain have been limited, with only one other suspected outbreak in the Philippines in 2014 [2]. NiV_M_ has an average CFR of ∼46%, a clinical presentation that is predominantly neurologic/encephalitic, and human-to-human transmission is limited [1, 2]. NiV strain Bangladesh (NiV_B_) has been responsible for recurrent outbreaks in Bangladesh and India since 2001 and has higher CFR than NiV_M_ at ∼75% [3–8]. Clinical disease is also different, with NiV_B_ patients presenting with both respiratory and neurologic signs [3–8]. Additionally, human-to-human transmission rates of NiV_B_ are higher compared to NiV_M_ [9, 10].

Pathogenic differences have been seen in animal models of NiV_M_ and NiV_B_. In African Green Monkeys (AGMs), 100% of the animals exposed to NiV_B_ succumb to disease by Day 7 post-exposure (PE), and these animals developed a more severe respiratory disease compared to NiV_M_ exposed animals [11]. Disease characteristics of NiV_M_ in AGMs are more variable [11–13]. Approximately 58% of animals succumb due to acute phase disease, typically between Days 9-12 PE, with respiratory signs the predominant finding. Animals that survive past the acute phase may develop a later stage encephalitic disease, resulting in severe neurological deficits often necessitating euthanasia [12, 13].

In hamsters exposed to NiV_M_ or NiV_B_ by the intraperitoneal (IP) route, differences in clinical disease signs were not significant, with both respiratory and neurologic signs noted for animals regardless of strain used [14]. However, progression to a terminal state was more rapid for NiV_M_ animals compared to NiV_B_ animals, and the dose of virus resulting in death of 50% of the animals (LD_50_) was higher for NiV_B_ (528 50% tissue culture infectious dose [TCID_50_]) compared to NiV_M_ (68 TCID_50_) [14].

For reported studies in hamsters, there is a great degree of variability for NiV_M_ in dose, survival, and time-to-death for both the IP and intranasal (IN) routes of exposure. Although a few studies in hamsters have been conducted with NiV_B_, the vast majority of model development and countermeasure evaluation work has used NiV_M_. Most of the work published using hamster models for NiV is from a single source/site, and very few studies have investigated the survival characteristics of NiV when plaque forming unit (pfu) assays are used to assess exposure dose as opposed to TCID_50_ assays. NiV_B_ represents the predominant circulating strain and is associated with a notably higher CFR. Therefore, it is critical to define the infection parameters in NiV_B_ animal model development so that these models can be more heavily employed in vaccine and treatment testing efforts.

Herein, we performed studies in Syrian hamsters assessing the dose response (in pfu) to NiV_B_ and NiV_M_ following both IP and IN exposure. IP doses of 10^5^, 10^6^, and 10^7^ pfu were 83-100% lethal regardless of strain used, but a delayed mean time-to-death (MTD) of ≥1 day was observed for NiV_B_ compared to NiV_M_ for these doses. NiV_B_ administered by the IN route was 60% lethal at the highest dose evaluated (10^6^ pfu), whereas 100% lethality was achieved when the same dose of NiV_M_ was evaluated by this route. Differences in clinical presentation for the strains were not noted.

Subsequent to model characterization efforts, we assessed the utility of these models for countermeasure evaluations by using Favipiravir (Favi). Subcutaneous (SC) administration of Favi has been shown to provide complete protection against lethal IP challenge with NiV when administered twice daily (BID) at a loading dose of 600 mg/kg/dose on the day of challenge, and a maintenance dose of 300 mg/kg/dose on Days 1-13 PE [15]. In the present study, two SC doses were evaluated on the same schedule as described by Dawes *et al* against both NiV_B_ and NiV_M_: 600/300 (loading/maintenance) mg/kg/dose and 300/150 mg/kg/dose. MTD increased for treated animals in the NiV_B_ groups regardless of dose, but similar results were not seen for NiV_M_ exposed animals, and there was no significant difference in percent survival or viral load for any treated groups compared to strain-matched controls. Contrary to the findings by Dawes *et al*, the results suggest that Favi as a treatment for NiV may not provide a significant protective advantage in the Syrian hamster model.

## Materials and methods

### Animals and housing

Research was conducted under an Institutional Animal Care and Use Committee (IACUC) approved protocol in compliance with the Animal Welfare Act, PHS Policy, and other Federal statutes and regulations relating to animals and experiments involving animals. The facility where this research was conducted is accredited by AAALAC International and adheres to the principles stated in the *Guide for the Care and Use of Laboratory Animals, National Research Council, 2011*. All efforts were made to minimize pain and distress.

One hundred and fourteen female Syrian golden hamsters (SGHs) (*Mesocricetus auratus*) aged 4-8 weeks at time of virus exposure, were obtained from Charles River Laboratories or Envigo for use on the dose ranging studies. Table 1 shows the study group designations, origins, and ages of the animals used on those studies. For the Favi efficacy study, forty-eight SGHs aged 4-6 weeks at time of virus exposure, were obtained from Envigo. Table 2 shows the study group designations for that study. There are no known pathogens endemic in the USAMRIID hamster colony. Housing rooms were maintained on a 12:12-h light:dark cycle (lights on at 0600 h, lights off at 1800 h) with fluorescent lights. Humidity was controlled between 30-70% and temperature was controlled between 68-76°F (set point 74.5°F). They were provided filtered (5 micron) domestic water via water bottles ad libitum. Water from animal rooms is tested quarterly (DoD Food Analysis and Diagnostic Laboratory, San Antonio, TX) for heavy metals, chlorinated hydrocarbons, nitrates, and microbial contamination. Hamsters were provided a species-appropriate Teklad Global diet (Harlan Teklad), fruits (as appropriate), and water *ad libitum* via an automatic watering system. Enrichment was provided as recommended by the *Guide for the Care and Use of Laboratory Animals*.

**Table 1.**
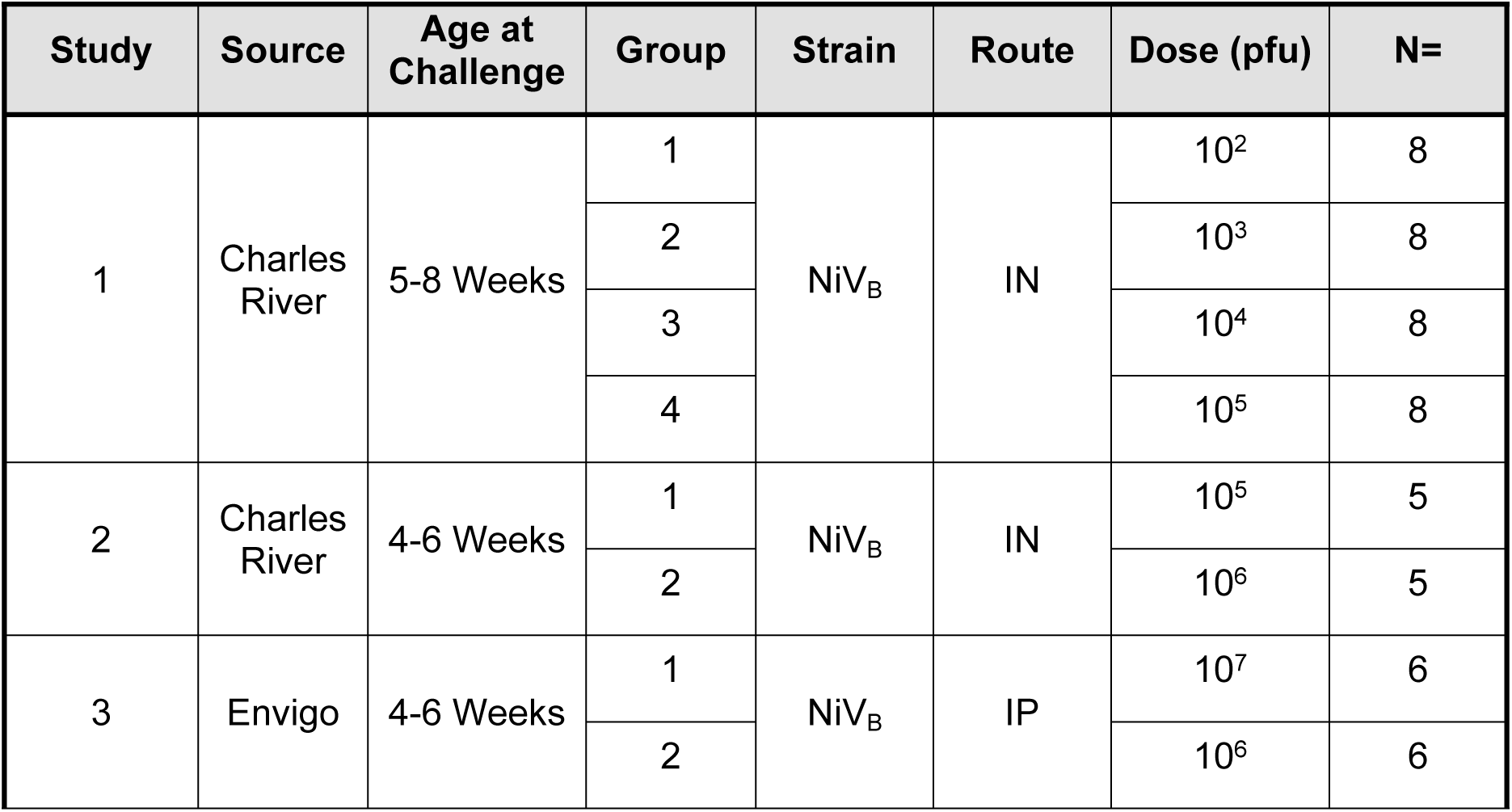
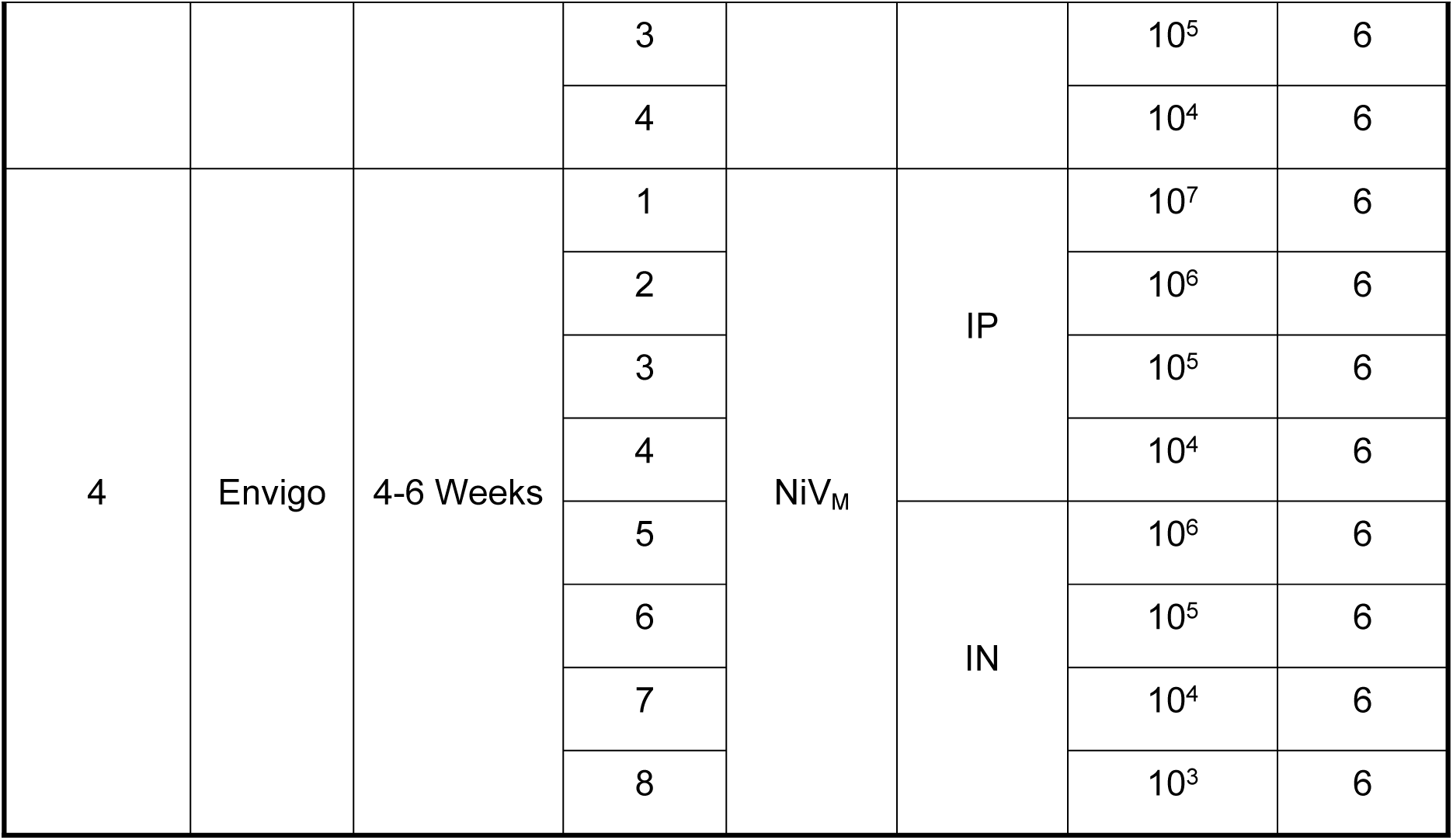
Study Group Designations – Dose Ranging Studies.

**Table 2.**
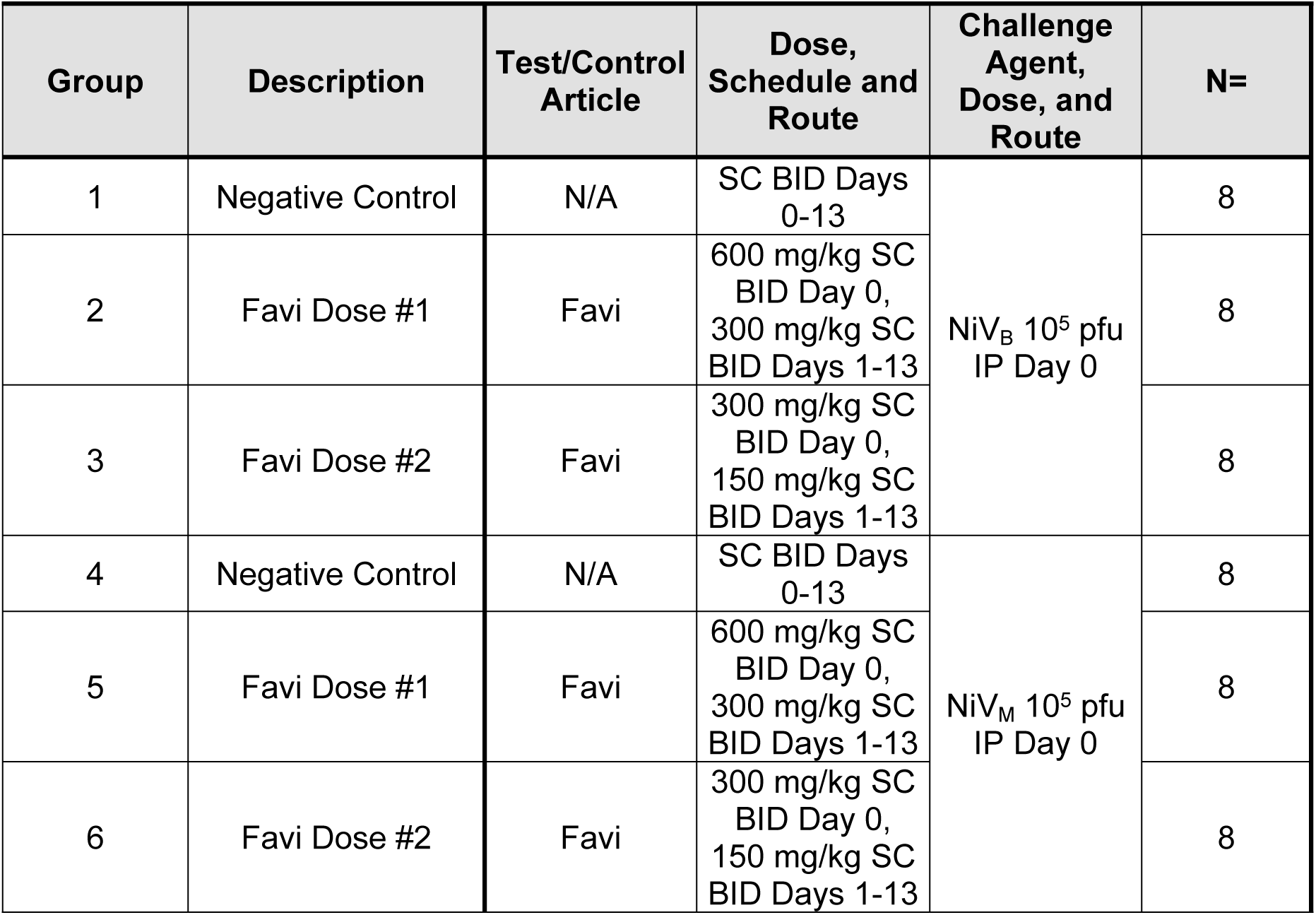
Study Group Designations – Favi Efficacy Study.

### Animal observations

SGHs were observed and scored daily in accordance with Table 3, beginning on the day of virus exposure (Day 0 PE). Observations were performed once daily while animals were healthy (score = 0), and up to three times daily, at appropriately spaced intervals, when clinical signs of illness were evident (score ≥1). Body weights were obtained once daily. SGHs that scored ≥8 met criteria to be considered moribund and were euthanized. Animals that survived to end of study were euthanized on Day 17 PE (dose ranging studies) or Day 21 PE (Favi study). Euthanasia was performed under deep anesthesia by barbiturate overdose in accordance with current *American Veterinary Medical Association Guidelines for the Euthanasia of Animals 2020* and institute standard operating procedures.

**Table 3.**
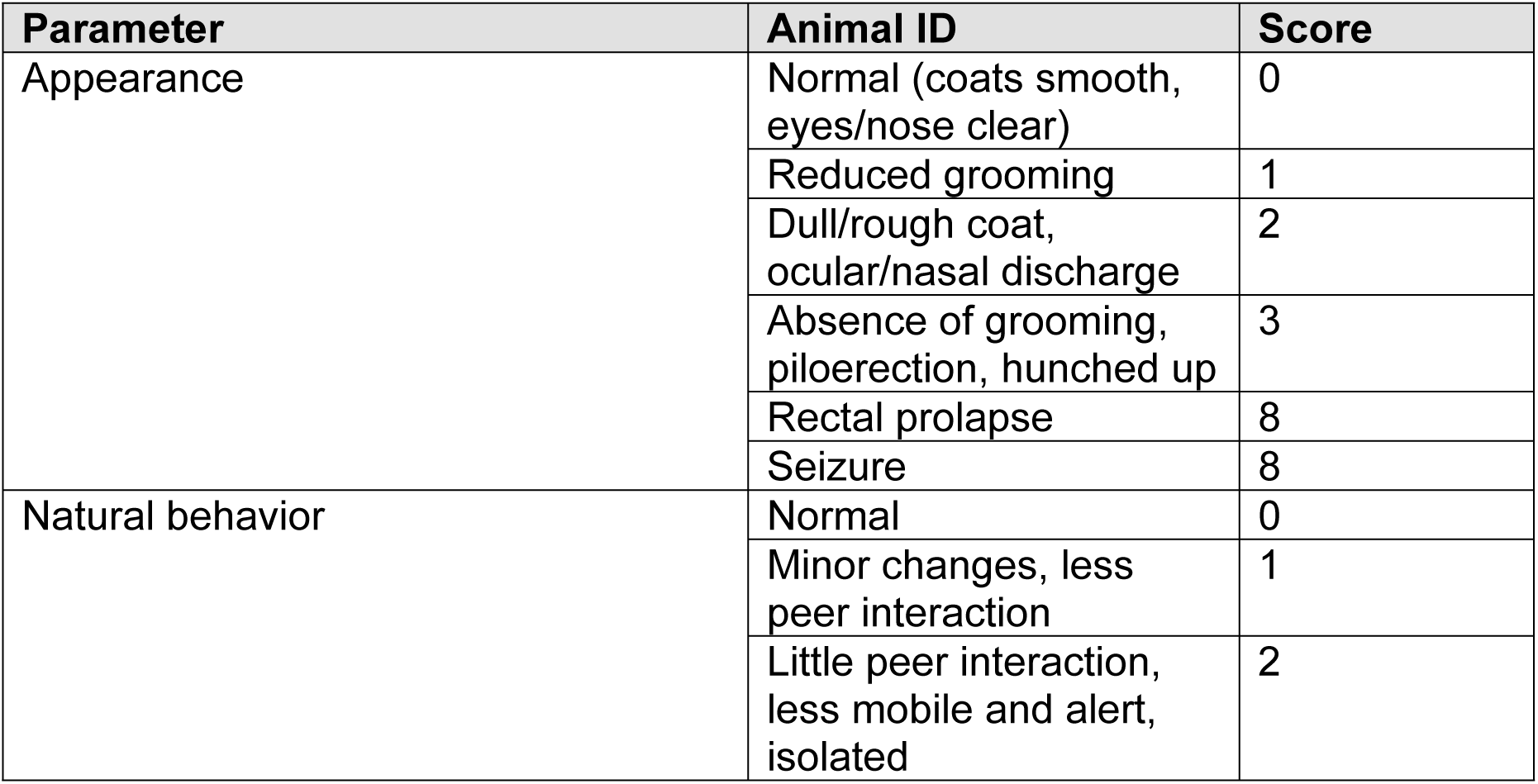
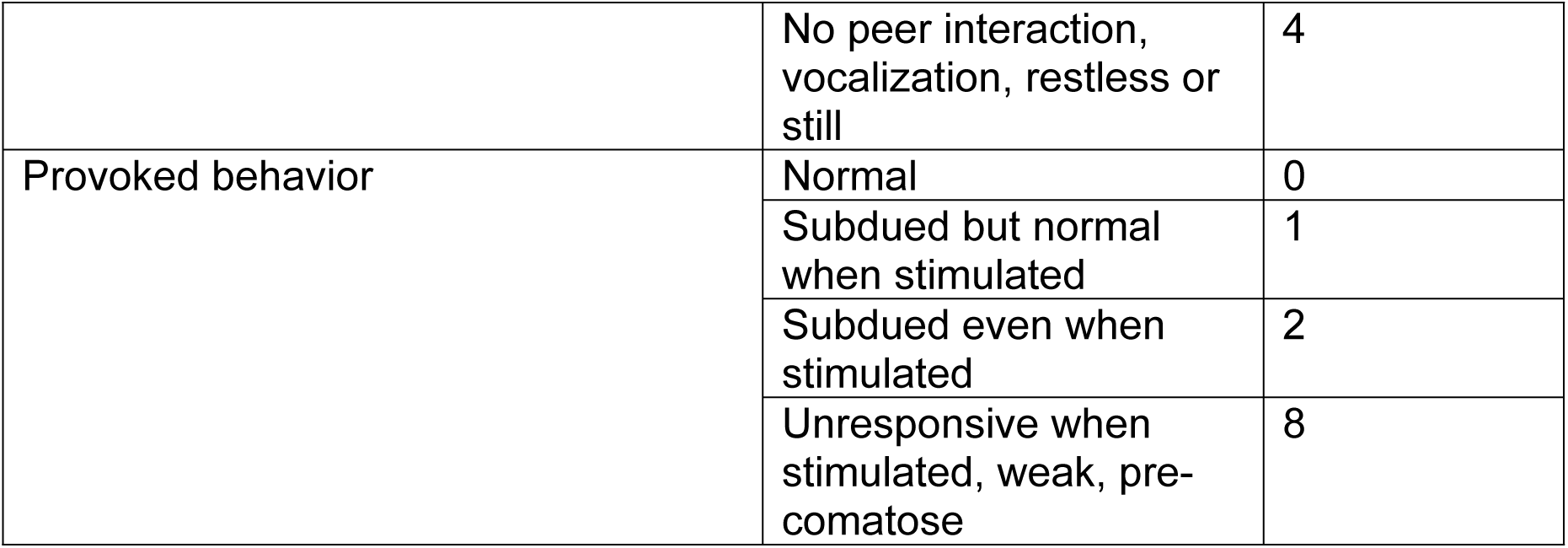
Clinical Signs Operational Definitions.

### Virus and virus exposure

NiV_B_ and NiV_M_ were obtained from Dr. Thomas Ksiazek at the University of Texas Medical Branch and were a first passage of the Centers for Disease Control and Prevention’s (CDC) isolate from the throat swab of SPBLOG# 200401006, patient ID# 3001 (2004). Research-grade seed stocks were produced at the United States Army Medical Research Institute of Infectious Diseases (USAMRIID) on Vero 76 cells (ATCC, Manassas, VA). These stocks were determined to be free of contaminants (bacterial, mycoplasmal) that could interfere with *in vitro* and *in vivo* experiments by bacterial culture plating and deep sequencing. Identity of the stocks were also verified by sequencing analysis.

On the day of virus exposure (Day 0 PE) animals were exposed either IP or IN to the target doses NiV_B_ or NiV_M_ in accordance with Tables 1-2. For IN inoculations, the dose was split equally into each nare, and total volume per nare was approximately 50µL. A sample of the exposure material was titrated by neutral red plaque assay to determine the injected pfu.

### Favipiravir preparation and administration

Favipiravir for use on study was formulated in water at a final stock concentration of 60 mg/mL after adding meglumine (Sigma Aldrich, Catalog # M9179) at a ratio of 1:1.2. Favi was then diluted in water to make a 30 and 15 mg/ml formulation. Syringe loading volumes were based on the average size of a 4-6 week old SGH of ∼100 grams, to achieve the desired mg/kg concentrations as shown in Table 2. The loaded volume per animal was ∼1 mL. Favi was administered SC BID on Days 0-13 PE.

### Blood Collection and Processing

Blood collection was only performed on the Favi study. Six animals per group were randomly chosen for blood collection on Days 0, 6, and 21 PE. However, for any of these selected animals that succumbed prior to the planned blood collection event on Day 6 PE, other animals in the same study group were substituted to assure to the extent feasible that samples for qRT-PCR and plaque assay were available for six animals in each group. For end-of-study collections, the same set of animals from Day 6 PE were used, if available; substitutions did not occur for any that had succumbed. Blood collection did not occur on hamsters found deceased. TRIzol™ LS (Invitrogen, Waltham, MA) was mixed with EDTA whole blood for qRT-PCR at a 3:1 ratio (3 parts TRIzol™ LS to 1 part EDTA whole blood) prior to RNA extraction and analysis.

### Extraction of viral RNA and quantification by qRT-PCR

Following the manufacturer’s procedure for QIAGEN’s QIAamp Viral RNA Mini Kit (250) (QIAGEN, Hilden, Germany), 70 µL of whole blood in TRIzol™ LS was manually extracted to produce a final purified elution volume of 70 µL. Purified nucleic acid was run in triplicate on the QuantStudio Dx platform (Applied Biosystems, Waltham, MA) using TaqPath 1-step RT-qPCR master mix (Applied Biosystems, Waltham, MA) according to the CDC’s recommended cycling conditions of 25°C for 2 minutes, 50°C for 15 minutes, and 95°C for 2 minutes, followed by 45 cycles of 95°C for 3 seconds and 60°C for 30 seconds. Primer and probe sequences, designed for NiV_B_, are as follows:

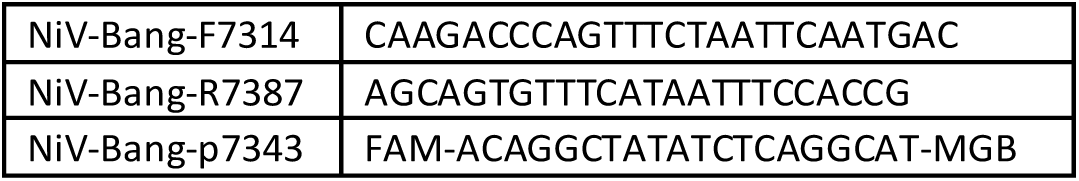

As a positive extraction control (PEC), 70 µL of the NiV_B_ stock in TRIzol™ LS at the previously-described 3:1 ratio was extracted alongside study samples to validate extraction efficacy. The PEC must produce viable PCR data across all three replicates. As a negative extraction control (NEC), 70 µL of nuclease-free water was extracted alongside study samples. All NEC replicates must be undetected. Nuclease-free water was run in triplicate and processed alongside study samples as a negative template control (NTC) to validate PCR efficacy. The NTC must be undetected. To create a standard curve, 70µL of the TRIzol™ LS containing stock virus was extracted, and a 1:10 serial dilution was performed down to dilution factor five. This is used to ensure precision and accuracy while quantifying the concentration of unknown samples. The standard curve must produce an R^2^ value greater than or equal to 0.99.

At least two out of three detected replicates were required to call a study sample positive. Samples for which one or no replicates were detected were considered negative.

### Plaque assay

Vero 76 cells (ATCC, Manassas, VA) at 85-100% confluency were used for this plaque assay, and 10-fold serial dilutions of study samples were prepared in plaque assay media (Minimum Essential Medium [MEM] [Corning, Manassas, VA] + 5% FBS [Corning, Manassas, VA] + 1% Penicillin/Streptomycin [x100] [Corning, Manassas, VA]). Dilutions were plated in triplicate, with 100 µL added to each well of a 6-well plate.

Plates were incubated at 37±2°C, 5±1% CO_2_ for 60±10 min, rocking every 15±5 min. Following incubation, 2 mL of primary agarose overlay (1% agarose [Lonza, Walkersville, MD] mixed 1:1 with 2x EBME (Gibco, Waltham, MA) was added to each well, and plates were incubated at 37±2°C, 5±1% CO_2_ for 3 days. After 3 days, 2 mL of secondary agarose overlay (1% agarose mixed 1:1 with 2x EBME and containing 4% neutral red (Sigma Aldrich, St. Louis, MO) was added to each well, and plates were incubated at 37±2°C, 5±1% CO_2_ overnight. Plaques were counted and virus titer was calculated based on all dilution series with 10-150 countable plaques. If wells at two different serial dilutions presented countable plaques within this range, then the counts from wells with the more concentrated dilution were chosen for calculating titer.

### Statistical analysis

The survival rates were compared by Fisher’s exact test, and the times to death were analyzed by the Wilcoxon rank-sum test for the pairwise comparison between the groups. LD_50_ calculations were performed by the Reed-Muench method. Survival curves were constructed using the Kaplan-Meier method and compared using log-rank tests. The following statistical analyses were performed only on the Favi study. The Kruskal Wallis test was used for overall group comparisons of maximum intervention scores and maximum qRT-PCR values, and Wilcoxon rank-sum test was used for pairwise comparisons between groups where the Kruskal Wallis test resulted in a p value of ≤0.10. For survival, qRT-PCR, and maximum intervention score analyses, step-up Hochberg adjustments were applied for multiple pairwise comparisons. Repeated measures ANOVA was used to test effects of group, time from challenge, baseline observation, and the interaction of group and time on weight and temperature. ANOVA was also used to examine differences in weight and temperature between groups at each post-challenge day. Post-hoc Dunnett’s tests were used to examine pairwise differences between each group at each post-challenge day where ANOVA resulted in a p value of ≤0.10. Analyses were performed using SAS Version 9.4 (SAS Institute, Inc., Raleigh NC).

## RESULTS

### NiV inoculation by the IP route causes severe disease/death in Syrian golden hamsters

Forty-eight SGHs were randomized into eight study groups, each containing six animals (**Table 4**). Groups 1-4 received NiV_B_ IP at a target dose of 10^7^, 10^6^, 10^5^, or 10^4^ pfu

**Table 4.**
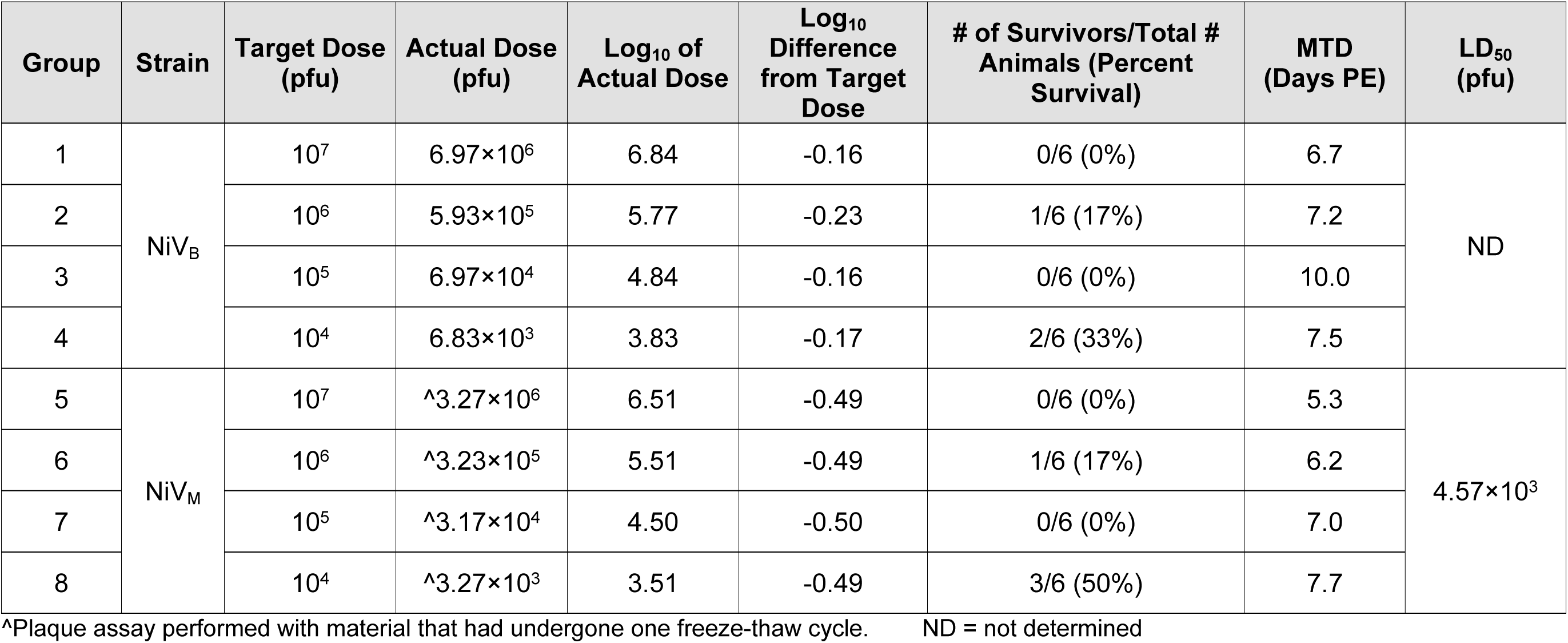
Outcome following IP inoculation with NiV.

(actual doses were 6.97×10^6^, 5.93×10^5^, 6.97×10^4^, and 6.83×10^3^ pfu, respectively) on Day 0 post-exposure (PE). Groups 5-8 received NiV_M_ at the same target doses (actual doses were 3.27×10^6^, 3.23×10^5^, 3.17×10^4^, and 3.27×10^3^ pfu, respectively) on Day 0 PE. SGHs were observed daily for signs of disease and were euthanized when moribund. Animals that survived until end-of-study were euthanized on Day 17 PE. Regardless of strain, all animals in the 10^7^ and 10^5^ pfu groups either succumbed or were euthanized due to terminal illness, with only one animal at 10^6^ pfu for both NiV_B_ and NiV_M_ surviving until end-of-study (**Fig 1**). Statistically significant differences in percent survival and MTD were not noted between dose groups, regardless of strain, and an LD_50_ was only able to be determined for NiV_M_ (4.57×10^3^ pfu). Survival time was significantly different (p<0.05) when the NiV_M_ 10^4^ pfu dose group was compared to the 10^5^ and 10^7^ pfu dose groups, because 50% of the animals in this dose group survived until end-of-study.

**Fig 1.**
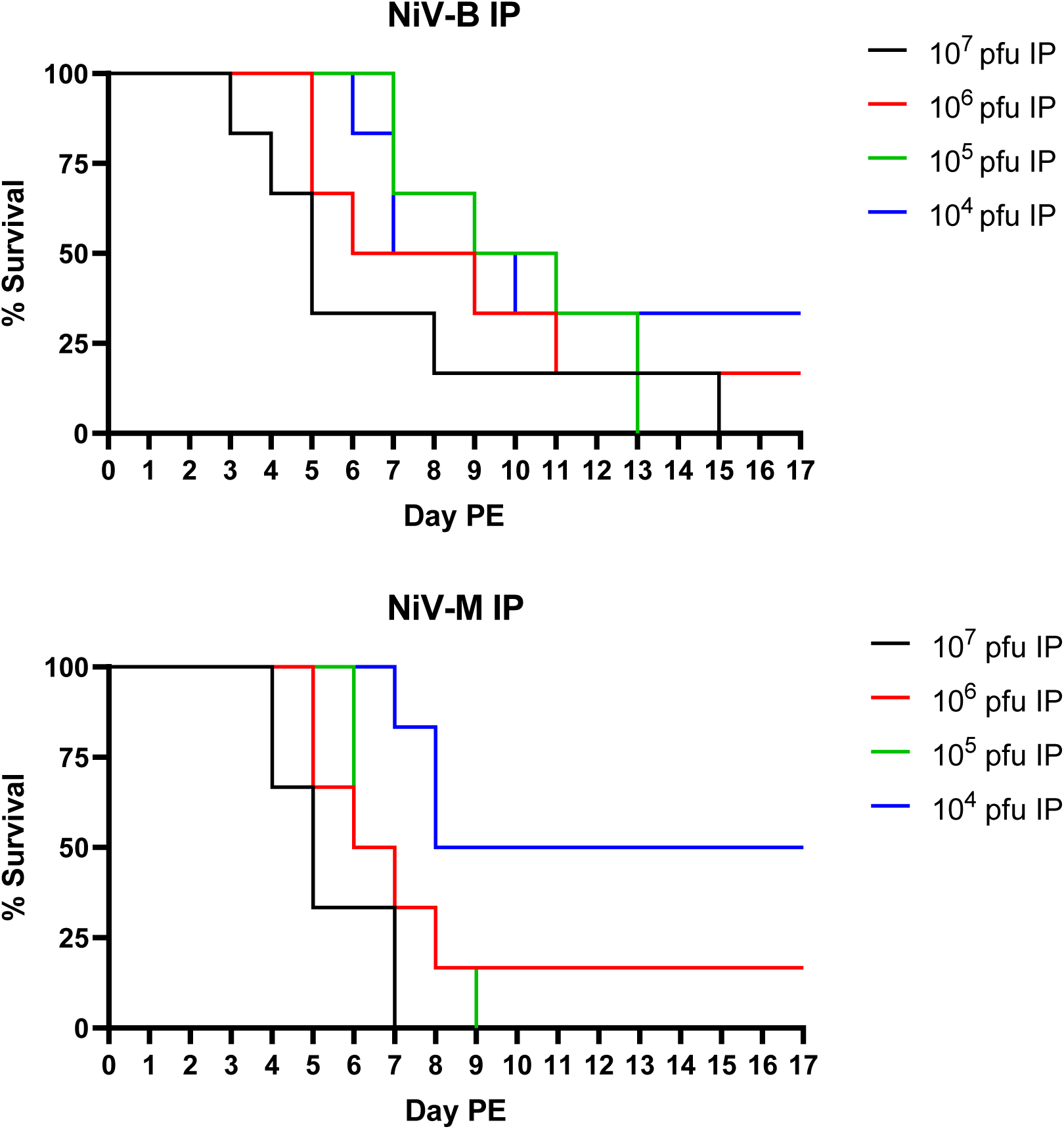
Survival following IP exposure to NiV_B_ or NiV_M_. Percent survival is shown for NiV_B_ (top panel) and NiV_M_ (bottom panel) for each of the target dose groups assessed. Clinical signs of disease for non-survivors were consistent across virus strains and exposure doses, and included an absence of grooming, piloerection, hunching, reduced responsiveness or unresponsive, rash, and neurological signs (head tilt, circling, balance issues, tremors, and/or seizures). Clinical signs were far less severe for survivors. When present, they included decreased grooming, nasal discharge, and/or a more subdued behavior; rash, tremors, and head tilt were noted for a few animals. Weight loss and fever were not prominent findings for this model, regardless of virus strain and exposure dose.

### High doses of NiV are required to cause clinical disease following IN inoculation of Syrian golden hamsters

Sixty-six SGHs were randomized into nine study groups (**Table 5**). Groups 1-5 (containing 5, 13, 8, 8, and 8 SGHs, respectively) received NiV_B_ IN at a target dose of 10^6^, 10^5^, 10^4^, 10^3^, or 10^2^ pfu (actual doses were 5.77×10^5^, 1.75×10^5^, 2.43×10^4^, 2.03×10^3^, and 2.83×10^2^ pfu respectively) on Day 0 PE. Groups 6-9 received NiV_M_ at a target dose of 10^6^, 10^5^, 10^4^, or 10^3^ pfu (actual doses were 3.27×10^5^, 3.23×10^4^, 3.17×10^3^, and 3.27×10^2^ pfu, respectively) on Day 0 PE. SGHs were observed daily for signs of disease and were euthanized when moribund. Animals that survived until end-of-study were euthanized on Day 17 PE. The only IN dose that resulted in uniform lethality was 10^6^ pfu of NiV_M_, and the MTD of 9.3 days was three days longer than the matched dose by the IP route (6.2 days) (**Fig 2**). The MTD of 11.0 days for NiV_B_ at 10^6^ pfu was nearly two days longer compared to the matched dose of NiV_M_ (9.3 days), and was nearly four days longer compared to the matched IP dose (7.2 days). In addition, 40% of the animals in this group survived until end-of-study. Less than or equal to 33% lethality was noted for all other dose groups for both strains, and no significant differences in MTD for non-survivors between dose groups was noted, regardless of strain. There were significant (p<0.05) differences in percent survival and survival time when the 10^6^ pfu dose groups were compared to other dose groups (exceptions: percent survival for NiV_M_ 10^6^ and 10^5^ pfu dose groups, and NiV_B_ 10^6^ and 10^2^ pfu dose groups, was not significantly different). As mentioned for the IP route, differences in survival time can be attributed to the large number of survivors in most dose groups. The LD_50_ for NiV_B_ by IN inoculation was 3.74×10^5^ pfu, and for NiV_M_ was 4.32×10^4^ pfu. Clinical signs of disease for survivors and non-survivors were consistent with those described following IP inoculation.

**Fig 2.**
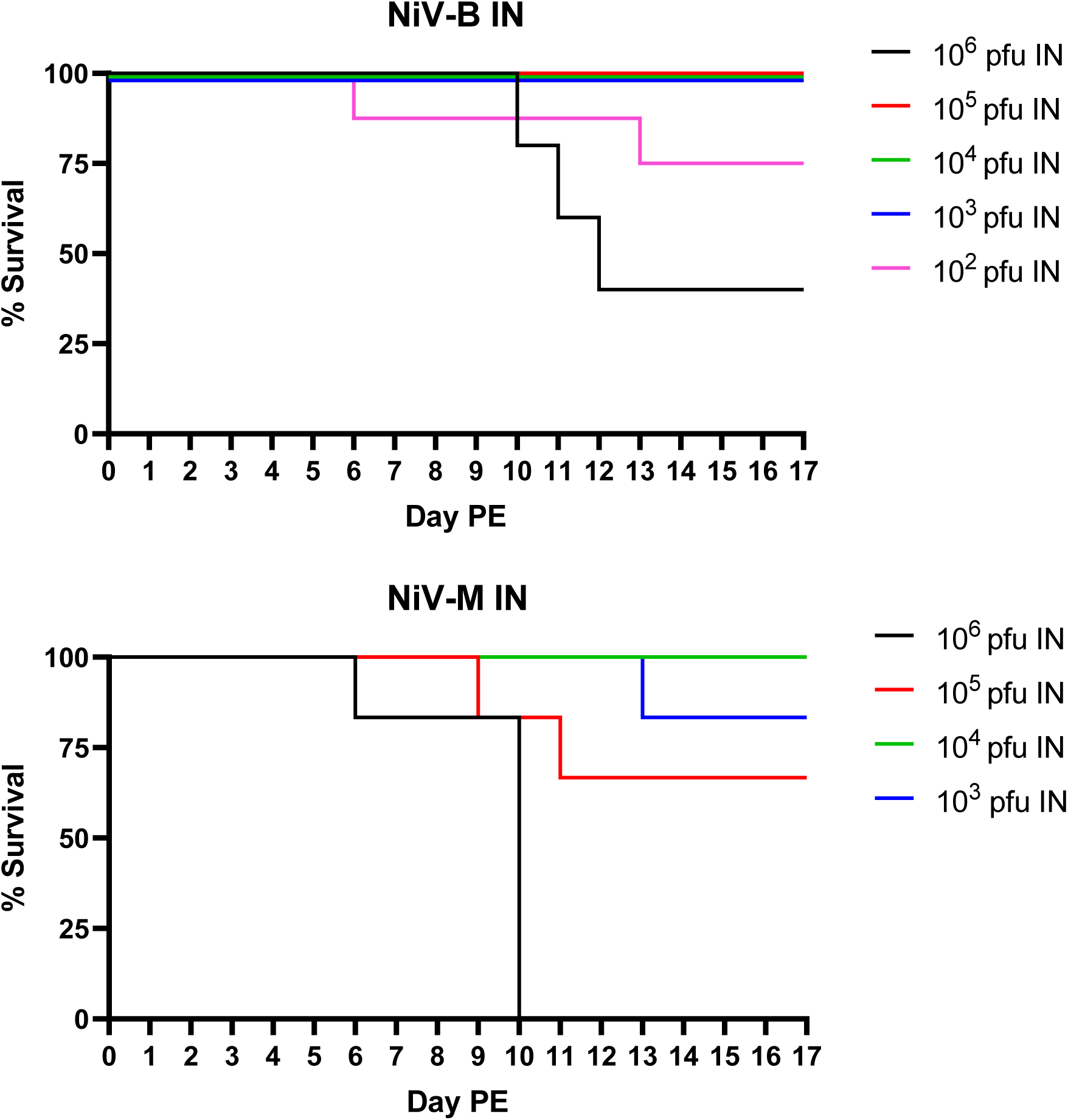
Survival following IN exposure to NiV_B_ or NiV_M_. Percent survival is shown for NiV_B_ (top panel) and NiV_M_ (bottom panel) for each of the target dose groups assessed.

**Table 5.**
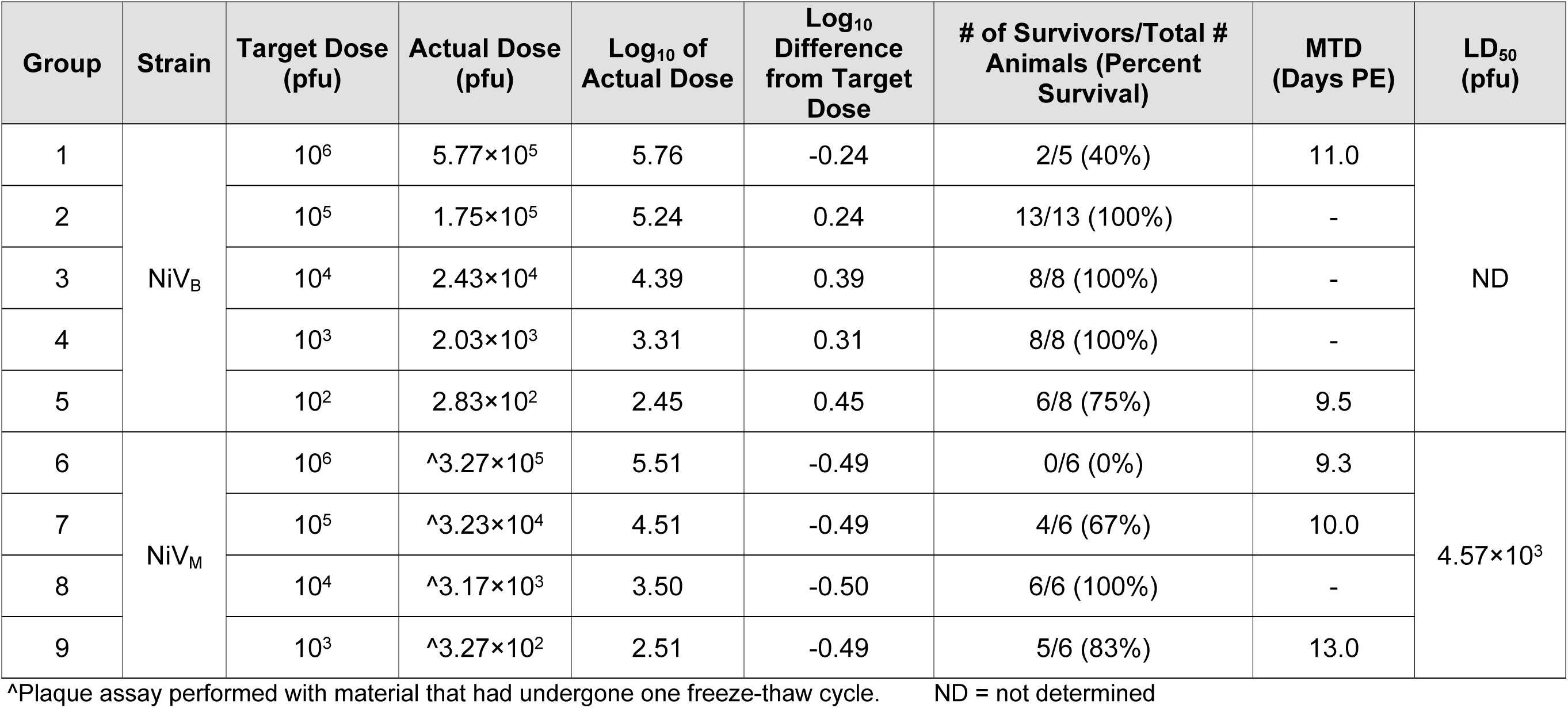
Outcome following IN inoculation with NiV.

### Favi is partially protective against severe disease/death following IP exposure of Syrian golden hamsters to NiV

Model characterization efforts described above suggested that the IP model is the most consistent at providing a survival endpoint that is desirable for countermeasure evaluations for both strains of NiV. This model was subsequently used to evaluate the efficacy of Favi against NiV_B_ and NiV_M_. Fourty-eight SGHs were randomized into six study groups, each containing eight animals (**Table 6**). Groups 2 and 5 received Favi at a 600/300 mg/kg (loading/maintenance) dose, and Groups 3 and 6 received Favi at a 300/150 mg/kg dose. Treatments were BID SC on Days 0-13 PE. On Day 0 PE, animals were inoculated IP with a target dose of 10^6^ pfu of either NiV_B_ (Groups 1-3) or NiV_M_ (Groups 4-6). SGHs were observed daily for signs of disease and were euthanized when moribund. Animals that survived until end-of-study were euthanized on Day 21 PE. All but one control animal per virus strain succumbed or was euthanized due to terminal disease. Fifty percent survival was obtained for both 600/300 mg/kg treatment groups (Groups 2 and 5) and the NiV_B_-exposed 300/150 mg/kg treatment group (Group 3) (**Fig 3**); survival dropped to 25% for the NiV_M_-exposed 300/150 mg/kg treatment group. Percent survival was not significantly higher than controls for both doses evaluated, regardless of virus strain. However, MTD was significantly longer (p<0.05) for Favi treated and NiV_B_ challenged animals, regardless of Favi dose.

**Fig 3.**
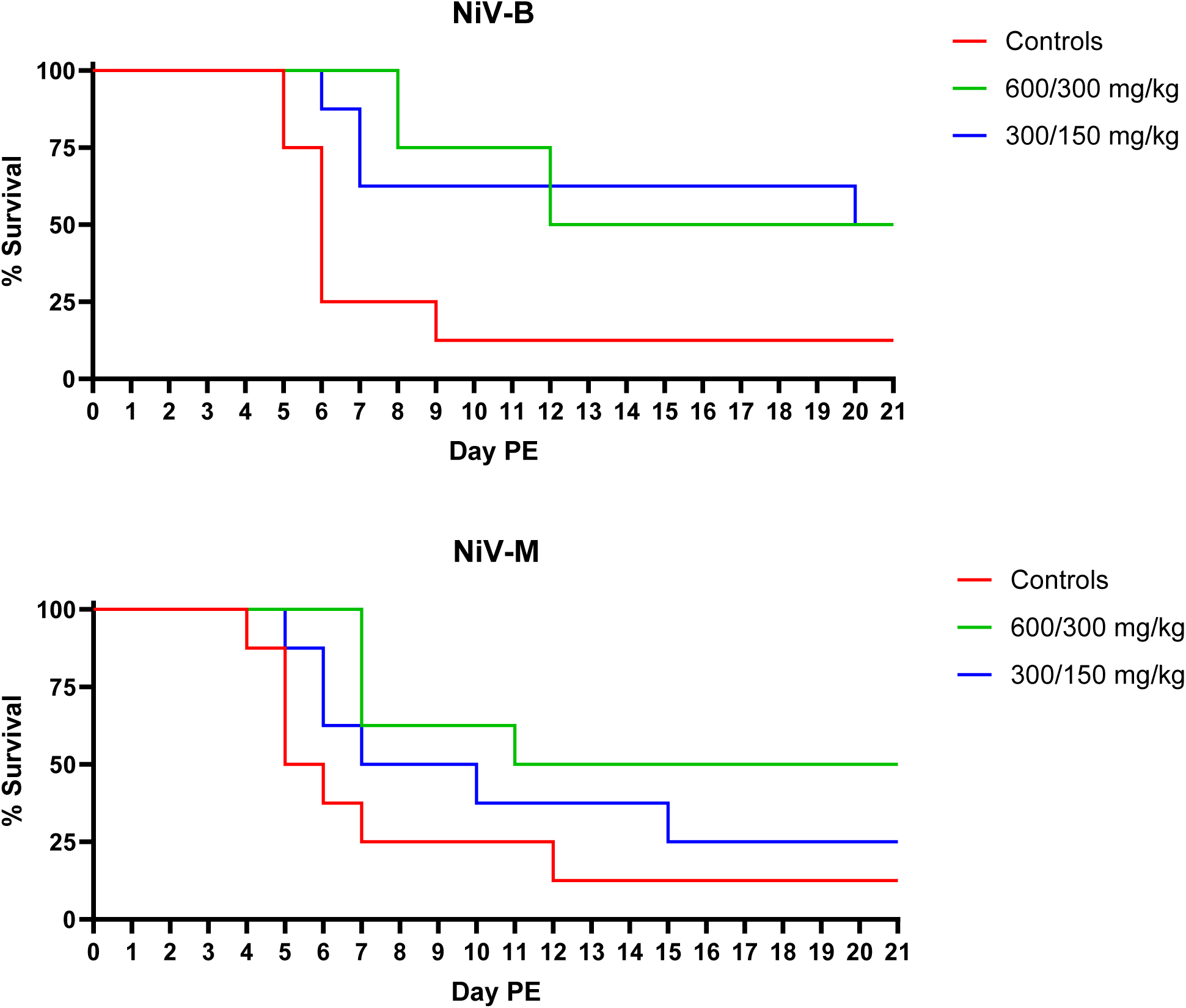
Survival following Favi treatment and IP exposure to NiV_B_ or NiV_M_. Percent survival is shown for NiV_B_ (top panel) and NiV_M_ (bottom panel) for each of the treatment doses assessed.

**Table 6.**
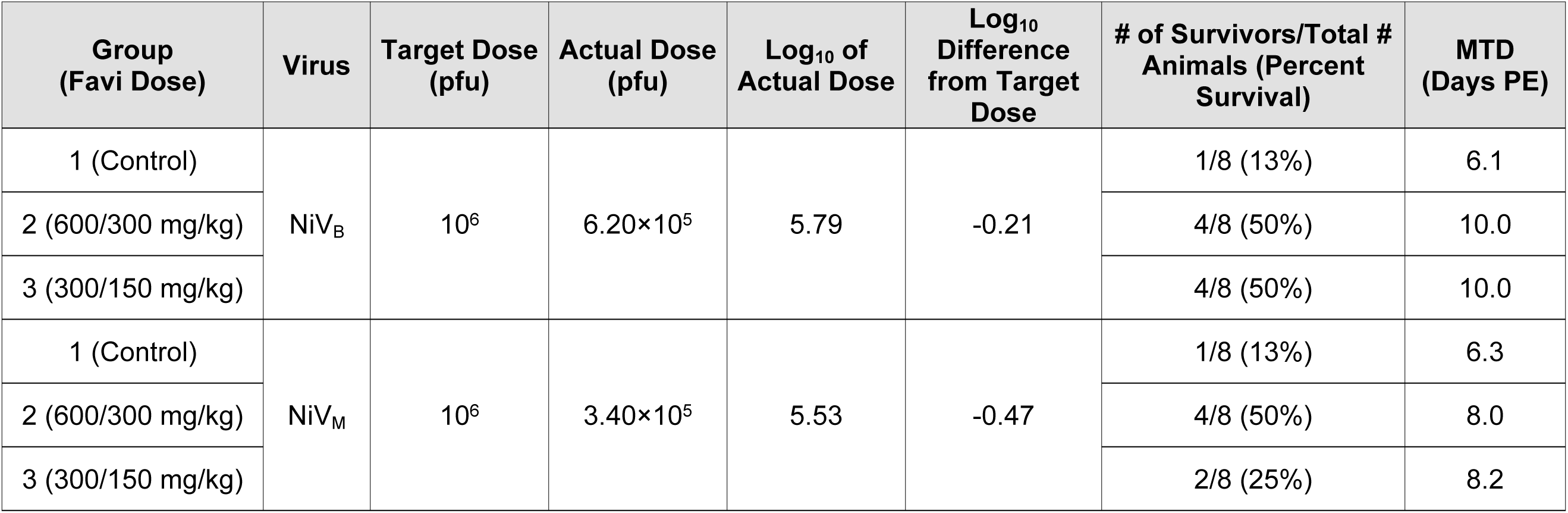
Outcome following Favi treatment and IP inoculation with NiV.

Clinical signs of disease for both survivors and non-survivors were as previously described in the model characterization studies.

Intervention scores for each animal were determined from daily observations from Day 1 PE through Day 21 PE (**Fig 4**). Animals were scored based on three parameters, as described in **Table 3**: appearance, natural behavior, and provoked behavior. Animals that were found dead were given an intervention score of 12 based on the maximum score for natural and provoked behavior. A notable 2-3 day delay in clinical disease, based on intervention scoring (**Fig 4**), was evident for 600/300 mg/kg treatment groups compared to strain matched controls. The 12-24 hour delay in disease for 300/150 mg/kg groups reveals a dose down affect but does not indicate a significant delay in disease presentation compared to controls. However, the maximum intervention score for Group 3 (300/150 mg/kg + NiV_B_) was significantly (p<0.05) lower than Group 1 (NiV_B_ controls).

**Fig 4.**
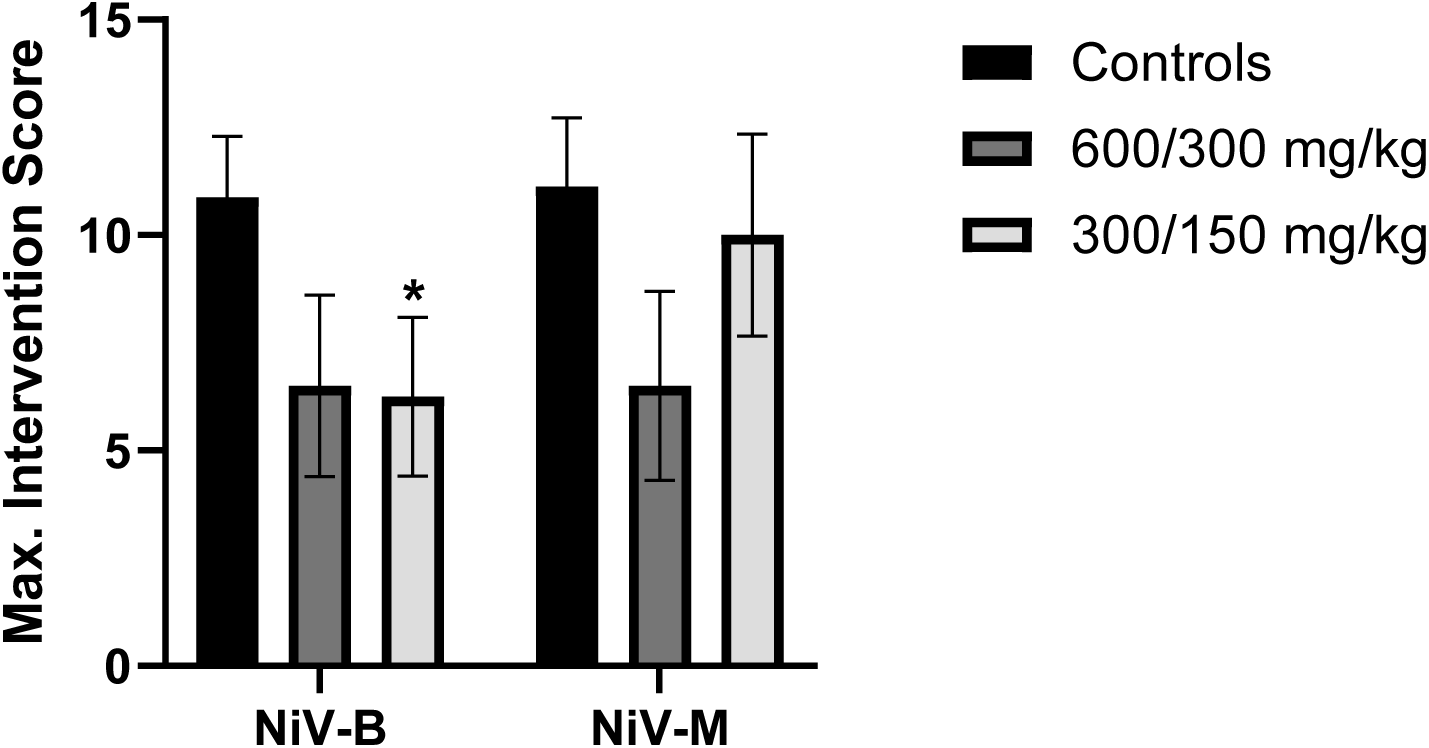
Group average maximum intervention scores. Bars represent the group mean, with error bars representing the standard error of the mean (SEM). Significant differences (p<0.05) between treatment groups and controls are indicated with an *.

Viral load in the blood was assessed by RT-qPCR for up to six animals in each study group on Days 0, 6, and 21 (end-of-study) PE. The majority of the animals in control groups, as well as the NiV_M_ treatment groups had viral RNA in blood on Day 6 PE, versus approximately 17% of the NiV_B_ treatment groups animals on the same day. With the exception of the 600/300 mg/kg treatment groups, the presence of viral RNA in blood by Day 21 PE was limited to the surviving NiV_M_ vehicle animal and one animal in each of the 300/150 mg/kg treatment groups. For both NiV_B_ and NiV_M_, all remaining animals in the 600/300 mg/kg groups assessed at Day 21 PE had measurable levels of viral RNA in their blood. Peak RNA levels were 1-2 Log_10_ copies/mL higher for controls compared to treated animals. However, this difference was not statistically significant.

## Discussion

In agreement with previously published studies [14, 16–18], exposure of SGHs to NiV by the IP route resulted in a consistent clinical disease characterized by both behavioral and neurologic findings. Lethality of greater than or equal to 50% was obtained with doses as low as 10^4^ pfu, and 80-100% lethality was obtained with doses as low at 10^5^ pfu regardless of NiV strain used, Malaysia or Bangladesh. The MTD was noticeably shorter for the 10^7^ pfu groups compared to other groups, regardless of strain. The relatively short MTD for this dose is not ideal for therapeutic evaluations as the window of time from exposure and/or clinical disease onset to death may be too truncated to accurately determine a protective effect. Consistent with previous efforts [14, 18], significant differences in MTD and survival time between NiV_B_ and NiV_M_ were observed when the 10^5^ pfu dose groups were compared (p=0.0476).

In general, the IN route of NiV exposure was far less consistent in terms of lethality than the IP route, and unlike the IP route, there was a clear performance difference between NiV_M_ and NiV_B_. For NiV_B_, 10^6^ pfu resulted in only 60% lethality. For NiV_M_, uniform lethality occurred at the 10^6^ pfu dose, but survival was greater than 60% for all other groups. In addition, the MTD for the 10^6^ pfu dose groups was 3-4 days longer compared to the matched IP dose groups, and this difference was statistically significant for NiV_M_ (p=0.0108). The data are consistent with previous reports [14, 16, 19] and suggest that the IN route may not be ideal for countermeasure studies where a consistent survival endpoint and/or NiV strain comparisons are required. If used, group sizes may need to be increased to ensure adequate statistical power is maintained.

It was previously reported that oral or SC administration of Favi 600/300 mg/kg (loading/maintenance) BID for 14 days (beginning on the day of virus exposure) was completely protective against NiV_M_ disease and death in hamsters [15]. To confirm these results and determine if Favi is a viable positive control for NiV efficacy studies, two doses of Favi, 600/300 mg/kg (loading/maintenance) and 300/150 mg/kg, were administered SC to hamsters BID for 14 days, beginning on the day of exposure to either NiV_B_ or NiV_M_. In contrast to published results, suboptimal protection was afforded by Favi, with only fifty percent of the animals surviving in the highest dose groups, regardless of virus strain. One possible explanation is the dose of virus used. In the Dawes *et al* study, an IP dose of 10^4^ pfu of NiV_M_ resulted in uniform lethality of vehicle control animals by Day 7 PE. However, in the model characterization studies discussed herein an IP dose of 10^4^ pfu was not uniformly lethal, with thirty-three percent survival for NiV_B_ exposed animals, and fifty percent survival for NiV_M_ exposed animals. The Favi positive control verification study was conducted using a dose of 10^6^ pfu. This dose was chosen because 10^4^ pfu was not uniformly lethal, and 10^5^ pfu of NiV_B_ had a significantly extended MTD compared to all other dose groups. A dose of 10^6^ pfu provided for near uniform lethality with a MTD that was consistent with other dose groups and between strains. However, at this virus dose, Favi was not effective at preventing disease or death in hamsters. It is, therefore, possible that a dose of 10^4^ pfu NiV by the IP route was too low to accurately predict overall Favi efficacy, but a dose of 10^6^ pfu IP created an environment that was too stringent due to virus overload. Collectively, the data suggest that further studies should be conducted to determine the effect of virus dose on Favi efficacy in the NiV hamster model.

## Acknowledgements

The use of either trade or manufacturers’ names in this report does not constitute an official endorsement of any commercial products. This report may not be cited for purposes of advertisement. Opinions, interpretations, conclusions, and recommendations are those of the author and are not necessarily endorsed by the U.S. Army or Department of Defense. Funding for this effort was provided by the National Institutes of Allergy and Infectious Diseases under Intra-agency agreement number AAI21016-001-00000.

^ Team Chenega, Contractor – this does not constitute an endorsement by the US Government of this or any other contractor.

† Laulima Government Solutions, Contractor – this does not constitute an endorsement by the US Government of this or any other contractor.

## Notes

### Competing Interest Statement

The authors have declared no competing interest.

